# Efficient and accurate inference of microbial trajectories from longitudinal count data

**DOI:** 10.1101/2020.01.10.902163

**Authors:** Tyler A. Joseph, Amey P. Pasarkar, Itsik Pe’er

## Abstract

The recently completed second phase of the Human Microbiome Project has highlighted the relationship between dynamic changes in the microbiome and disease, motivating new microbiome study designs based on longitudinal sampling. Yet, analysis of such data is hindered by presence of technical noise, high dimensionality, and data sparsity. To address these challenges, we propose LUMINATE (LongitUdinal Microbiome INference And zero deTEction), a fast and accurate method for inferring relative abundances from noisy read count data. We demonstrate on synthetic data that LUMINATE is orders of magnitude faster than current approaches, with better or similar accuracy. This translates to feasibility of analyzing data at the requisite dimensionality for current studies. We further show that LUMINATE can accurately distinguish biological zeros, when a taxon is absent from the community, from technical zeros, when a taxon is below the detection threshold. We conclude by demonstrating the utility of LUMINATE for downstream analysis by using estimates of latent relative abundances to fit the parameters of a dynamical system, leading to more accurate predictions of community dynamics.

**Code availability:** https://github.com/tyjo/luminate

## 1 Introduction

The human body is home to trillions of microbial cells that play an essential role in health and disease ^5^. The gut microbiome, for instance, encodes over 3 million genes ^20^ responsible for a variety of normal physiological processes such as the regulation of immune response and breakdown of xenobiotics^6^. Disturbances in gut communities have been associated with several diseases, notably obesity^17^ and colitis ^18^, and changes to the vaginal microbiome during pregnancy is associated with risk of preterm birth ^7^. Consequently, investigating the human microbiome can provide insight into biological processes and the etiology of disease.

A major paradigm for microbiome studies design uses targeted amplicon sequencing of the 16S rRNA gene to produce read counts of each bacterial taxon in a sample^16^. Due to its low cost (compared to shotgun metagenomics), 16S rDNA sequencing is a valuable tool for generating coarse-grained profiles of microbial communities. Nonetheless, analysis of 16S datasets faces multiple domain-specific challenges. First, 16S datasets are inherently compositional ^12^: they only contain information about the relative proportions of taxa in a sample. In addition, technical noise, such as uneven amplification during PCR, can produce read counts that differ substantially from the underlying community structure ^16^. In particular, species near the detection threshold may fail to appear in a sample, necessitating a distinction between a biological zero — where a species is absent in the community — from a technical zero where it drops below the detection threshold^1^. Finally, the number of taxa and time points in a sample may be large, requiring methods that scale to high dimensional data.

Increasingly, study designs based on 16S rDNA sequencing have incorporated longitudinal sampling. This is exemplified by a major aim of the second phase of the Human Microbiome Project ^19^ being quantification of dynamic changes in the microbiome across disease-specific cohorts. Longitudinal sampling holds promise in elucidating causality between temporal changes in the microbiome and disease. It further provides a unique opportunity to address the statistical challenges of 16S sequencing by pooling information across longitudinal samples.

To this end, two recent methods have been proposed for analyzing noisy longitudinal count data: TGP-CODA^1^ and MALLARDs^22^. TGP-CODA fits a Gaussian process model to longitudinal count data, providing estimates of denoised (latent) relative abundances and statistical correction for technical zeros. MALLARDs, dynamic linear models with multinomial observations, fit a state space model to count data to partition observed variation into biological and technical components. Both models highlight the importance of temporal modeling, and its utility in providing insight into microbial systems. However, efficient inference from time-series data is a challenging problem, and both methods have difficulty scaling with sample size and taxa.

### 1.1 Our contribution

We propose LUMINATE (LongitUdinal Microbiome INference And zero deTEction), an accurate and efficient method to infer relative abundances from microbial count data. Our contribution is two-fold. First, using variational inference we reformulate the problem of posterior inference in a state-space model as an optimization problem with special structure. Second, we propose a novel approach to differentiate between biological zeros and technical zeros.

We demonstrate on synthetic data that LUMINATE accurately reconstructs community trajectories orders of magnitude faster than current approaches. We further demonstrate LUMINATE’s ability to accurately distinguish biological zeros from technical zeros. Finally, we demonstrate the utility of LUMINATE by using estimated relative abundances to infer the parameters of a dynamical system, leading to more accurate predictions of community trajectories.

## 2 Methods

### 2.1 Probabilistic Model of Latent Variables

Methods for inference from time-series data are often formulated using state-space models. State-space models describe latent dynamics as a sequence of time-indexed random vectors, ***x***_*t*_, where ***x***_*t*_ is dependent on time points in the past. Information about the hidden state of the system is obtained through noisy observation of each time point ***y***_*t*_. Such models are well suited for describing microbial dynamics: ***x***_*t*_ contain information about the true — hidden — relative abundances, while ***y***_*t*_ are noisy sequencing reads. Furthermore, state-space models provide a flexible framework for more sophisticated modeling that better captures the data generating process. We include two additional variables important for modeling microbial count data: ***w***_*t*_ which describes extinction and recolonization of taxa, and ***z***_*t*_ which incorporates an additional layer of sequencing noise(Figure 1).

**Figure 1:**
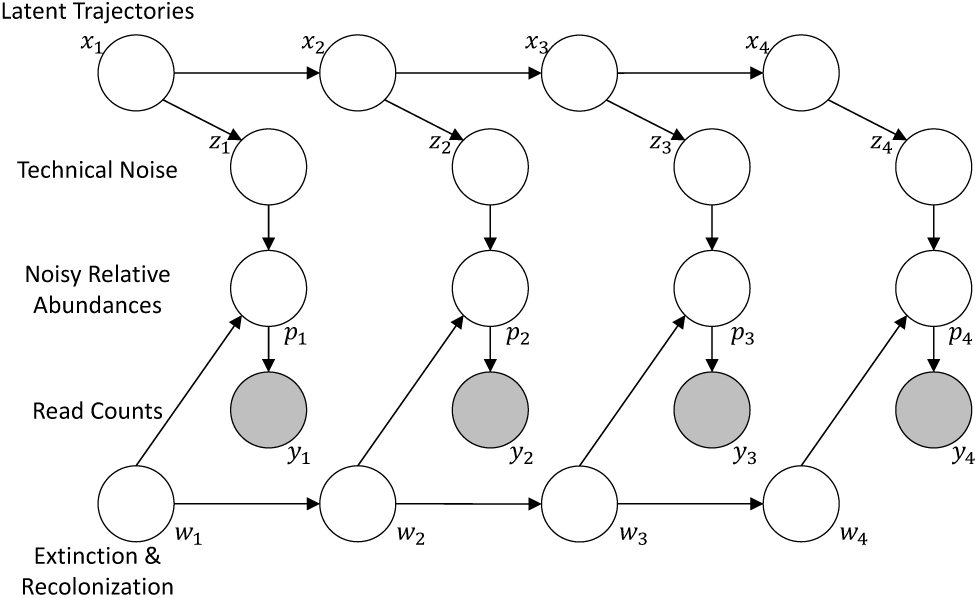
Graphical model for 4 time points. Sequencing counts ***y***_*t*_ are determined by noisy relative abundances ***p***_*t*_, which themselves are determined by the taxa alive at time *t*, ***w***_*t*_, and noisy realizations ***z***_*t*_ of the true community state ***x***_*t*_.

Specifically, our model is as follows. Suppose we have a sample with *T* observed time points. Let 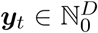 be the sequencing reads among *D* taxa at time *t*, and let ***x***^*t*^ ∈ ℝ^*D*-1^ be the additive log ratio of the relative abundances of those taxa (the natural parameters of the multinomial distribution). The time between observations *t* - 1 and *t* is denoted Δ_*t*_. Further, let ***z***_*t*_ ∈ ℝ^*D*-1^ be variables that represent noisy realizations of ***x***_*t*_, and let 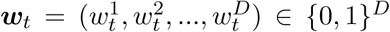 be indicator variables denoting which taxa are alive at time point *t* (i.e. 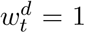 if taxa *d* is alive at time *t*, 0 otherwise). Our model is given by:

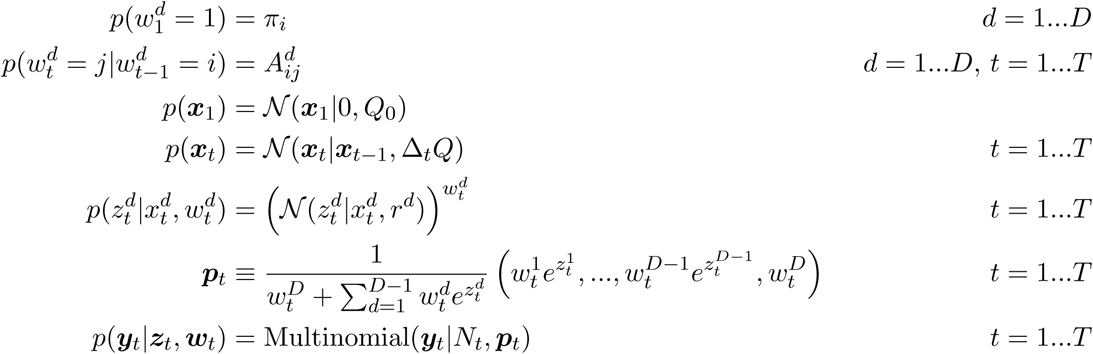

The ***z***_*t*_ describe additional sequencing noise not captured by the multinomial distribution. The multinomial distribution makes a strong assumption that the technical variance is purely due to otherwise uniform statistical sampling. The 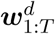 constitute a hidden Markov model with transition probabilities 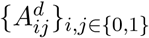 describing the extinction and reintroduction of certain taxa, distinguishing biological from technical zeros. Setting 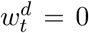 removes the contribution of 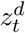 from the likelihood, and zeros out the relevant proportions in the multinomial counts. Conceptually the ***w***_1:*T*_ approximate extinction and recolonization events by making them orthogonal to the state of the system. Finally, the ***x***_*t*_ serve as a prior over the space of dynamics. The change in the system between time points depends on the covariance between ratios of taxa *Q*, and the time between observations Δ_*t*_. By learning the posterior ***x***_1:*T*_ |***y***_1:*T*_, we can estimate relative abundances from sequencing counts through ***x***_1:*T*_.

The covariance *Q* implicitly makes the assumptions that trajectories are smooth in time. However external perturbations such as antibiotics can rapidly induce changes in the community. We model these changes by introducing a perturbation covariance *Q*_*p*_ that replaces *Q* for time points with known (i.e. provided as input) perturbations.

Our model is conceptually similar to TGP-CODA^1^ and MALLARDs^22^. Both models introduce variables analogous to ***z***_*t*_ for technical noise, but take different approaches to modeling dynamics that come with increased computational cost. MALLARDs use a similar state-space model (that describes dynamics under a phylogenetically motivated log-ratio transformation). However, MALLARDs require evenly spaced time points — each time point occurs after a fixed interval of time. After specifying a unit of time, time points without observations are integrated out computationally using a Kalman filtering/smoothing approach. Additionally, MALLARDs do not incorporate terms for biological zeros as we do here. TGP-CODA, in contrast, incorporates additional variables for technical zeros similar to ***w***_*t*_, but not true zeros which we claim the ***w***_*t*_ represent. Furthermore, TGP-CODA learns a state-space covariance matrix using a Gaussian process model. This increased flexibility comes at a considerable computational burden.

### 2.2 Inference

Our main contribution is the demonstration that inference under such state-space models can be performed quickly using variational inference without loss of accuracy. By inference, we mean two things: posterior inference where the goal is to compute the posterior *p*(***x***_1:*T*_, ***z***_1:*T*_, ***w***_1:*T*_ |***y***_1:*T*_), and parameter inference for the model parameters *A*^1:*D*^, *Q*, and *r*^1:*D*-1^. Variational inference transforms both inference problems to an optimization problem by approximating the true posterior *p*_*θ*_(·|***y***) with model parameters *θ* by a variational posterior *q*_*ν*_ (·|***y***) with variational parameters *ν*. The parameters (*θ, ν*) are optimized to minimize the Kullback-Leibler divergence, or equivalently maximize the “evidence lower-bound”, between the true and approximate posterior. The variational objective function is

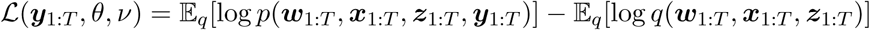

The main challenge in designing an inference algorithm for variational inference is choosing a form for *q* that is capable of closely approximating the true posterior while maintaining the ability to compute the expectations in ℒ (while black-box approaches exist where the expectations in ℒ are not explicitly computed, a closed form inference procedure is more desirable). Assuming a particular factorization of *q* and optimizing parameters using coordinate ascent, it is sometimes possible to compute an optimal parametric form for *q* for that also gives the optimal *ν* (see Blei *et al*. ^3^ for a derivation).

A common choice of factorization is to partition model variables into independent subsets

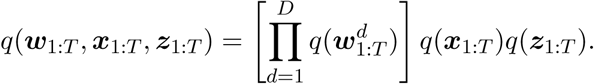

For this choice of factorization, the optimal 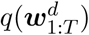 and *q*(***x***_1:*T*_) can be computed in closed form using block coordinate ascent (which we will show), while we need to make a choice for the parametric form of *q*(***z***_1:*T*_). A sensible choice for 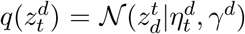. The 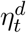 and *γ*^*d*^ are variational parameters that are optimized with respect to ℒ. The joint distribution across ***z***_1:*T*_ is

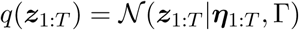

where Γ is a diagonal covariance matrix with entries in {*γ*^1^, …, *γ*^*D*-1^}. Given this choice of *q*(***z***_1:*T*_) the optimal choice of *q*’s for *q*(***x***_1:*T*_) and 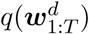 are given by ^3^

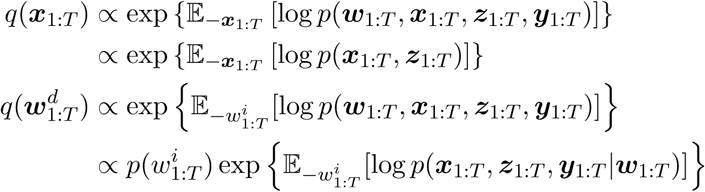

where the expectations are computed with respect to all *q* except for the variable of interest. We devote the remainder of this section to demonstrating that these can be computed efficiently in closed form.

First, the joint distribution of *p*(***x***_1:*T*_) = 𝒩(0, Λ^-1^) is Gaussian with precision matrix Λ that is block tridiagional. The simplest way to see this is to note that a Gaussian density is equivalent to its Laplace approximation. Hence, Λ is given by

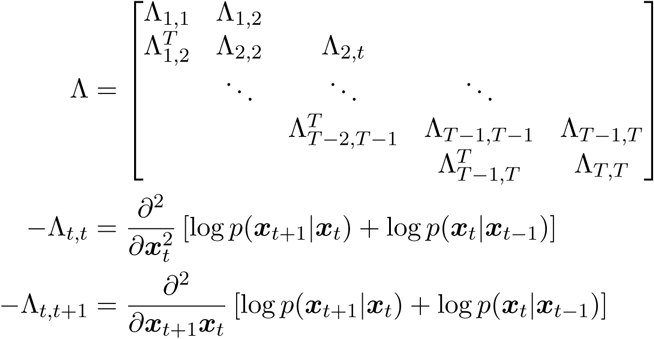

Simplifying 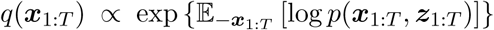leaves us with *q*(***x***_1:*T*_) = 𝒩(***x***_1:*T*_ |***µ***_1:*T*_, Σ) where Σ and ***µ***_1:*T*_ are given by

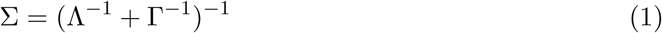

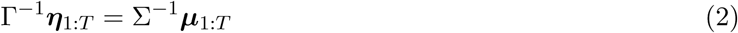

Notably, if we’re only interested the posterior means ***µ***_1:*T*_, we never need to explicitly compute the entire posterior covariance Σ. Σ^-1^ is block tridiagonal, which means its inverse can be computed in 𝒪(*TD*^3^) time instead of 𝒪(*T* ^3^*D*^3^) time ^13^. Furthermore, the solution for ***µ***_1:*T*_ only relies on the diagonal blocks of Σ^-1^ and an intermediate computation from the inverse. Consequentially, ***µ***_1:*T*_ can be computed in 𝒪(*TD*^2^) after the inverse is computed, instead of 𝒪(*T* ^2^*D*^2^).

Simplifying the expression for 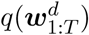, reveals that the optimal 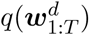is given by

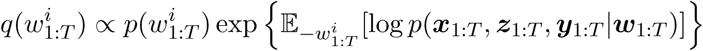

This is precisely the posterior under a hidden Markov model with (now fixed) observations given by the exponential term. Moreover, the only terms we need to compute ℒ are 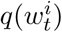 and 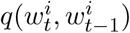, which can be computed in 𝒪(4*T*) time using the standard forward-backward equations for hidden Markov models (HMMs) ^2^.

Finally, the update for the parameters of *q*(***z***_1:*T*_) cannot be computed in closed form due to the structure of the problem. We instead rely on a conjugate gradient algorithm to optimize ***η***_1:*T*_ (since ***η***_1:*T*_ does not rely on the variance terms *γ*^*d*^ we choose not to optimize *γ*^*d*^).

The only remaining difficulty is computing

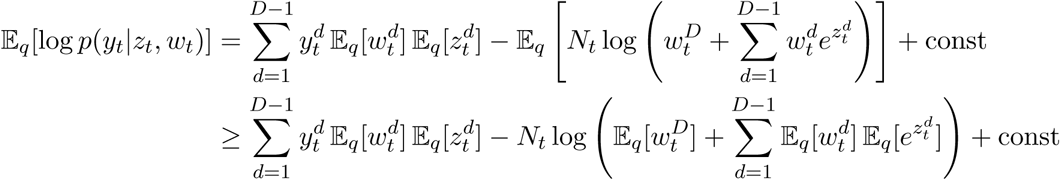

in ℒ. This lower bound on 𝔼_*q*_[log *p*(*y*_*t*_|*z*_*t*_, *w*_*t*_)] bounds the objective ℒ by below, which we note maintains a valid variational inference algorithm.

Once *q* has been formulated, optimizing model parameters *A*^1:*D*^, *Q, r*^1:*D*-1^ are straightforward. The expectations in L can all be computed (using the lower bound above), and taking the gradient with respect to each parameter and setting equal to zero obtain a closed form for each.

In summary, we have derived an inference algorithm for the model parameters and variational parameters of our model, where we can compute closed form block coordinate ascent updates for all but one set of parameters. Moreover, we can compute such updates efficiently by exploiting the special structure of the covariance of the state-space. Thus, we are left with the following algorithm.

#### Algorithm 1: LUMINATE’s inference algorithm

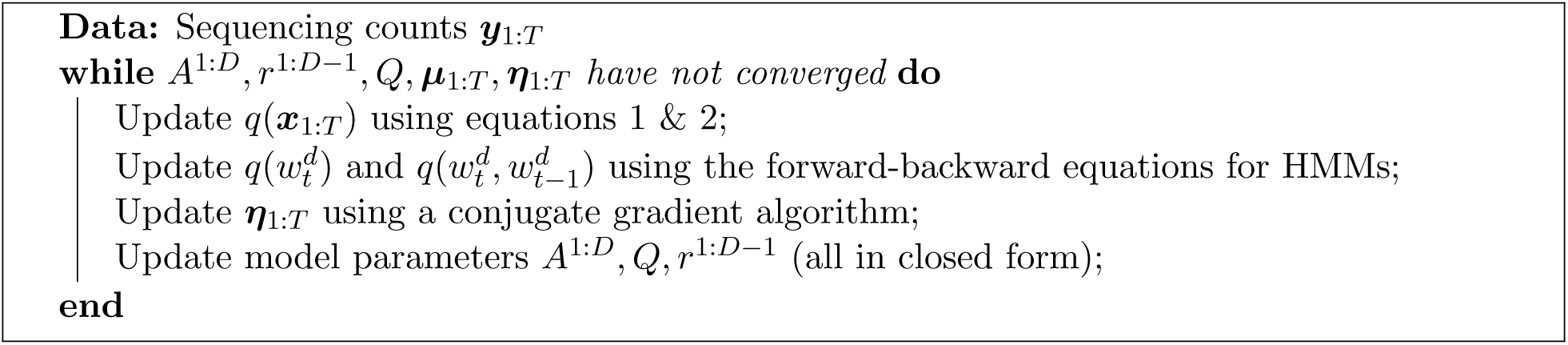

### 2.3 Simulation evaluation

We designed simulations to evaluate our model’s ability to infer relative abundances from noisy sequencing data. To this end, we downloaded two dense longitudinal datasets of bacterial concentrations from Bucci *et al.* ^4^: i) a dataset of 5 gnotobiotic mice colonized with a bacterial mixture of 16 species (the *C. diff* dataset), and ii) a dataset of 7 germ-free mice colonized with a mixture of 17 Clostridia strains (the *Diet* dataset). The *C. diff* dataset mice were subject to a *C. difficile* challenge after 28 days (average 26 observed time points observed over 56 days). The *Diet* dataset mice were fed a high-fiber diet for 5 weeks, switched to a low fiber diet for 2 weeks, then returned to the high-fiber diet for 2 weeks (average 47.14 observed time points across 65 days). We used these datasets to learn the parameters of a generalized Lotka-Volterra model (gLV) ^26^. We chose to simulate trajectories using gLV because gLV has been shown to accurately describe microbial dynamics in some cases, in particular on the datasets we used to generate model parameters (see Stein et al. ^26^ or Bucci et al. ^4^).

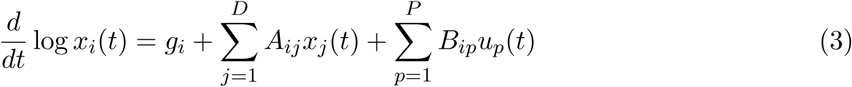

The *x*_*i*_(*t*) denote the concentration of bacteria *i* at time *t*, and the *u*_*p*_(*t*) denote external perturbations (such as introduction of *C. difficle* and change in diet). The parameters *g*_*i*_, *A*_*ij*_, and *B*_*ip*_ describe growth rates, interactions, and external effects respectively. We fit equation (3) by discretizing it and performing least squares with elastic net regularization, similar to Stein *et al.* ^26^.

Once we learned the model parameters for each dataset, we then forward simulated trajectories for each dataset given initial conditions of each mouse using the Runge-Kutta 5(4) method of numerical integration as implemented in `RK45` from `SciPy`^14^. This generated evenly spaced time points whose number corresponded to the number of observed time points of each mouse. We qualitatively inspected the simulated trajectories to ensure they matched the ground truth dynamics in the original data.

We simulated sequencing counts on top of each ground truth trajectory under varying levels of sequencing noise, following the framework of Silverman et al. ^22^. Briefly, given temporal covariance *Q* and noise covariance *R*, they defined a signal-to-noise ratio as the total variance of *Q* over the total variance of *R*

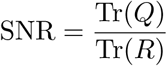

We computed the SNR under the additive log-ratio transformation: alr(*x*_*i*_(*t*)) = log (*x*_*i*_(*t*)*/x*_*D*_(*t*)), using alr(***x***(*t*)) = (alr(*x*_1_(*t*)), …, alr(*x*_*D*-1_(*t*)) to compute the a diagonal covariance matrix *Q* of the state-space give by alr(***x***(*t*)). The diagonal entries of *Q* measure how quickly each taxon alr(*x*_*i*_(*t*)) changes over time. For fixed *Q* and fixed SNR, we set *R* = diag{*r*_1_, …, *r*_*D*-1_} where 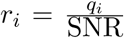. Thus, the sequencing noise was proportional to the variability of each taxa.

Finally, we simulated sequencing reads for each time point from the following model

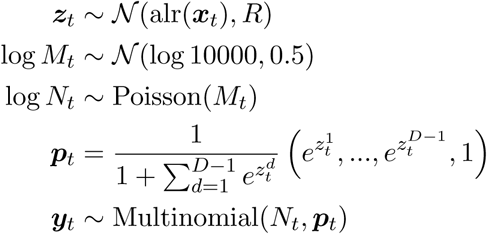

Intuitively, this means the average sequencing depth is approximately 10000 reads. The log-normal Poisson distribution on the number of sequencing reads *N*_*t*_ increases the variance in depth across samples, to better match the high variance of sequencing depth found in real data.

Importantly, all models we evaluated (see Section 2.5) make the same or more general assumptions about technical noise, and none assume gLV dynamics. Äijö et al. ^1^ assume a model equivalent to noise under the additive log-ratio transformation with additional noise from technical zeros, prior to observed sequencing counts. Silverman et al. ^22^ assume noisy realizations occur under the isometric log-ratio transformation (ilr). The ilr is a linear combination of the alr, and therefore simulating under the alr is equivalent to ilr under a linear transformation of the covariance matrix.

### 2.4 Biological zero detection simulations

To determine the ability of our model to detect biological zeros from technical zeros, we simulated 4 taxa across 30 days under gLV with carefully chosen parameters. We picked parameters such that one taxon would go extinct during the simulation, while forcing another taxon to remain near the detection threshold. The remaining 2 taxa were abundant throughout the simulation. This resulted in an approximately 2-to-1 ratio of true zeros versus technical zeros. We trained our model across 10 datasets of 10 longitudinal samples each, and for each observed zero computed the posterior probability that it was a biological zero: 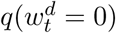.

### 2.5 Model comparison

We downloaded the code for TGP-CODA from GitHub ^11^. As TGP-CODA only runs on a single sample at once, we ran it on each sample in each dataset individually using the default parameters, then combined the results. We estimated latent relative abundances by taking the mean of the posterior samples of variables Θ_*G*_ computing using the No-U-Turn Sampler (NUTS) in `PyStan`^24^.

We downloaded the code for the MALLARD model from GitHub ^10^, and extracted the code that performed posterior inference under their model in `RStan`^25^. Because the MALLARD implementation is not a complete software package, we needed to perform two modifications to the code to run on our simulated data. First, we used the canonical basis instead of the phylogenetic basis for the isometric log-ratio transformation. This results in no loss of generality because it only affects the interpretation of the coordinates of the state-space. Second, we changed how samples for MCMC were initialized. The original implementation used `RStan`’s black box variational inference algorithm to compute initial samples before running the NUTS sampler. However, `RStan`’s black box variational inference can fail unexpectedly, so we resorted to initializing samples using `RStan`’s default initialization. We estimated relative abundances by transforming posterior samples of *θ* to relative abundances, then taking the mean.

### 2.6 Utility for downstream analysis

We used estimated relative abundances from LUMINATE to fit the parameters of “compositional” Lotka-Volterra (cLV) ^15^, a nonlinear dynamical system describing microbial relative abundances we recently proposed. cLV uses the following model to describe changes in relative abundances across *D* over time:

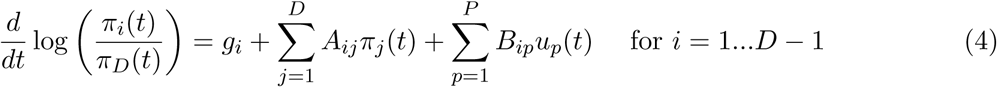

The *π*_*i*_(*t*) give the relative abundance of taxon *i* at time *t, g*_*i*_ its relative growth rate, *A*_*ij*_ the relative interactions between taxa, and *B*_*ip*_ the effect of external perturbations. We learned the parameters *A*_*ij*_, *B*_*ip*_, *g*_*i*_ by discretizing (4) and performing least squares with elastic net regularization, trained on the *π*_*i*_(*t*) estimated by LUMINATE. Regularization parameters were chosen using leave-one-out cross validation, picking parameters with the lowest prediction error from initial conditions.

We compared LUMINATE + cLV to two other time-series models: the sparse autoregressive model (sVAR) proposed by Gibbons et al. ^8^ and the ARIMA-Poisson model proposed by Ridenhour et al. ^21^. We download sVAR from GitHub ^9^, and ARIMA-Poisson from the supplementary material in Ridenhour et al. ^21^. We fit both models following the methods from each respective paper: ARIMA-Poisson was fit with 1 time lag, sVAR was fit with 3 time lags. We further rarefied OTU counts to 10,000 reads for sVAR following Gibbons et al. ^8^.

We compared model performance on two datasets by predicting trajectories from initial conditions on test data. The first dataset was the *C. diff* dataset described above (the *Diet* dataset included concentrations only). We also used a dataset of 6 white-throated woodrats fed oxalate from Ridenhour et al. ^21^. Using leave one-out cross validation, we predicted community trajectories on held out samples using model parameters learned on the remaining data.

## 3 Results

### 3.1 Simulations to assess the accuracy of LUMINATE

We first evaluated how well LUMINATE reconstructed (latent) community trajectories under varying amounts of sequencing noise. We generated ground truth trajectories by simulating data under generalized Lotka-Volterra (gLV) using parameters learned from real data (see Methods), then simulated noisy sequencing counts on top of each ground truth trajectory with varying signal-to-noise ratio and time between observations. We evaluated LUMINATE in comparison to three other models: i) a Dirichlet-Multinomial model (i.e a pseudocount model), ii) TGP-CODA^1^, and iii) the specific MALLARD model from Silverman et al. ^22^. Performance was compared by computing the mean *r*^2^ between true and estimated trajectory for each taxon across longitudinal samples. This beneficially treats rare and common taxa on an equal scale.

Encouraging, LUMINATE had a significantly higher *r*^2^ (p ¡ 0.05; Wilcoxon-signed rank test) than the Dirichlet-Multinomial model across all 8 of our simulations with evenly spaced time points (Figure 2A). We further observed significantly higher *r*^2^ in 6 of 8 simulations when compared with the MALLARD model, and on 7 out of 8 simulations when compared with TGP-CODA. Importantly, LUMINATE performed no worse than the competing models across any of the simulations we investigated. Taken altogether, this suggests that LUMINATE is better recreating the latent community dynamics.

**Figure 2:**
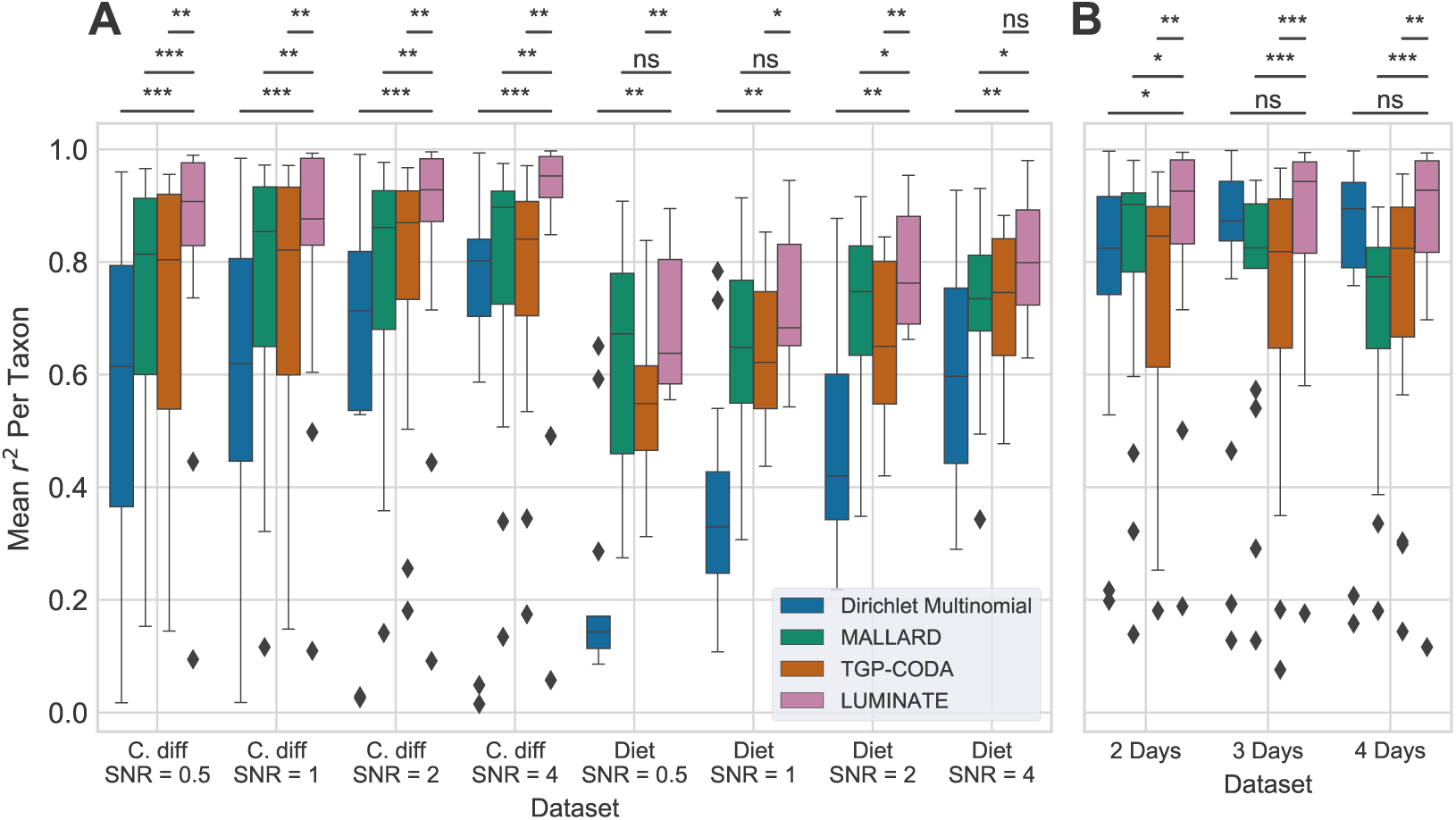
LUMINATE accurately recapitulates relative abundance trajectories. (A) Mean *r*^2^ (y-axis) between ground truth and estimated relative abundances trajectories for each taxon. Equally spaced time points were simulated under generalized Lotka-Volterra with parameters learned from two real datasets (*C. diff* and *Diet*) with varying signal-to-noise ratio (SNR; x-axis). (b) Effect of sampling time on estimated trajectories under the *Diet* simulations with SNR=4. (Wilcoxon signed-rank test; ns: not significant; *: *p <* 0.05; **: *p <* 0.01; ***: *p <* 0.001).

Real microbiome datasets tend to be sparse in time. We therefore performed simulations to investigate sensitivity to technical frequency. We simulated data under learned parameters from *C. diff* data, and removed time points so that there was an observation every 2, 3, and 4 days on average. Notably, LUMINATE was robust to the sparser simulations (Figure 2B), outperforming TGP-CODA and the MALLARD model on all three simulations.

### 3.2 Simulations to assess the efficiency of LUMINATE

Both TGP-CODA and MALLARD models rely on Markov Chain Monte Carlo (MCMC) algorithms to compute posterior estimates of model variables. As MCMC can be computationally expensive, we wanted to evaluate how each model scales with increasing number of observed time points and taxa. We thus simulated a single longitudinal sample varying the number of time points and taxa.

Across all datasets, LUMINATE was faster then the other methods we investigated (Figure 3), sometimes by more than 2 orders of magnitude. LUMINATE ran in *<* 1.5 minutes on all datasets. In contrast, it to the MALLARD model 8.3 hours to run 50 taxa at 10 time points. On this same dataset it took TGP-CODA 18.28 minutes to run, but 1.7 hours to run on 50 taxa at 30 time points. In practice, this means that LUMINATE is the only method that can scale to datasets with multiple longitudinal samples and many observed taxa.

**Figure 3:**
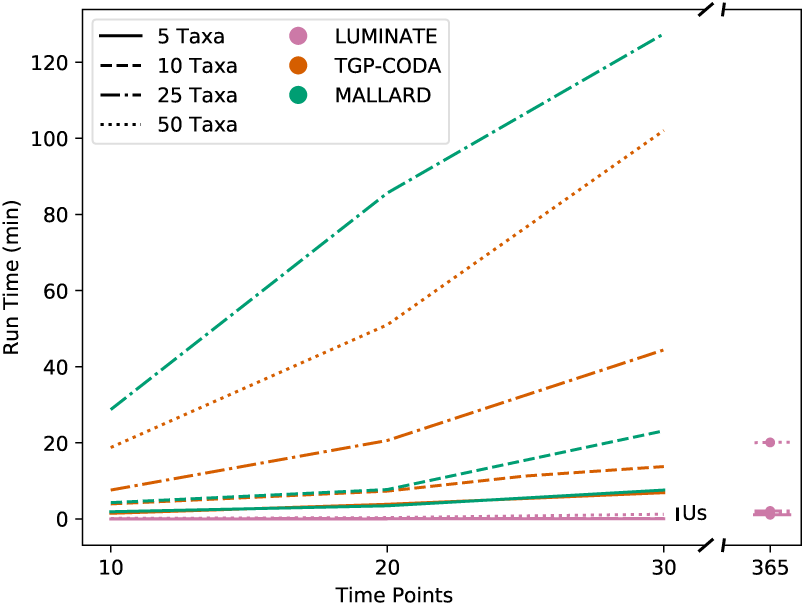
LUMINATE is efficient. Run time (measured as user time) in minutes (y-axis) for each model on a single longitudinal sample varying the number of time points (x-axis). Right: estimated run times for LUMI-NATE on 365 time points.

### 3.3 LUMINATE distinguishes biological zeros from technical zeros

We carefully designed simulations to test LUMINATE’s ability to distinguish biological zeros — where a taxon is not presenting the community — from technical zeros, where it is below the detection threshold. Specifically, we simulated data where one taxon goes extinct over the course of the simulation, while another hovers near the detection threshold. For all zeros in the observed data, we computed the posterior probability of a biological zero, and evaluated performance by computing the area under the receiver operating characteristic (AUC-ROC). This measures the probability of the event that a biological zero receives a higher posterior probability than a technical zero, an indicator that the model differentiates between the two. We performed 10 replicates with 10 samples each to estimate confidence intervals for the AUC-ROC. Notably, the mean AUC was high across all replicates (Figure 4; mean = 0.91, std = 0.04), suggesting that our model accurately discriminates biological from technical zeros.

**Figure 4:**
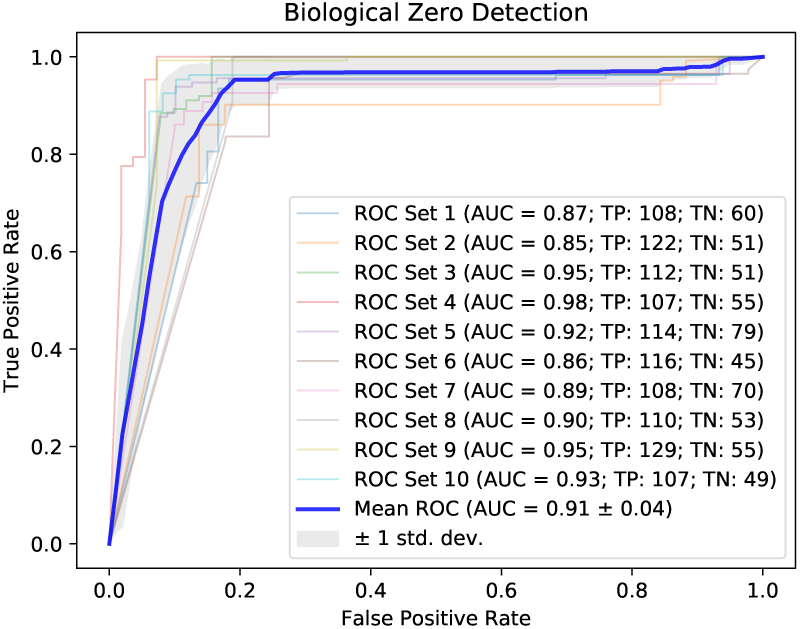
LUMINATE accurately discriminates biological from technical zeros. AUC-ROC curve using the posterior probability of a biological zero as a predictor for biological zeros on 10 simulated datasets. (TP: True Positives; TN: True Negatives)

### 3.4 Utility for downstream analysis

We have demonstrated that LUMINATE accurately estimates relative abundances. These “denoised” estimates can be useful for downstream analysis of longitudinal data. One example is learning the parameters of a dynamical system. We recently proposed a nonlinear dynamical system called “compositional” Lotka-Volterra (cLV) that describes how relative abundances change over time. However, learning the parameters of cLV requires estimated relative abundances (as would any other dynamical system describing relative abundances). We thus asked if LUMINATE could be useful for fitting nonlinear models of microbial dynamics, and if such a nonlinear model could lead to better descriptions of the underlying dynamics.

We fit cLV using LUMINATE’s estimated abundances, and used cLV to forecast community trajectories from initial conditions. We compared forecast trajectories to two other models: ARIMA-Poisson ^21^ and (sVAR) ^8^. For each model, we computed the average error between estimated and observed trajectories by taking Euclidean distance (the error) divided by the number of observed time points. We compared each model to LUMINATE + cLV by looking at the difference in error between the competing model and LUMINATE + cLV. This difference is expected to be symmetric around 0 if both models perform equally well (and greater than 0 if LUMINATE + cLV is performing better).

We observed that LUMINATE + cLV more accurately predicted trajectories than both sVAR and ARIMA-Poisson (Figure 5). For the sVAR model, this was significant in the Oxalate dataset. We observed a skew favoring LUMINATE + cLV on the *C. diff* dataset, but it did not reach the significance threshold, likely reflecting the smaller sample size (fewer taxa) in this dataset. In contrast, LUMINATE + cLV significantly outperformed ARIMA-Poisson on both datasets.

**Figure 5:**
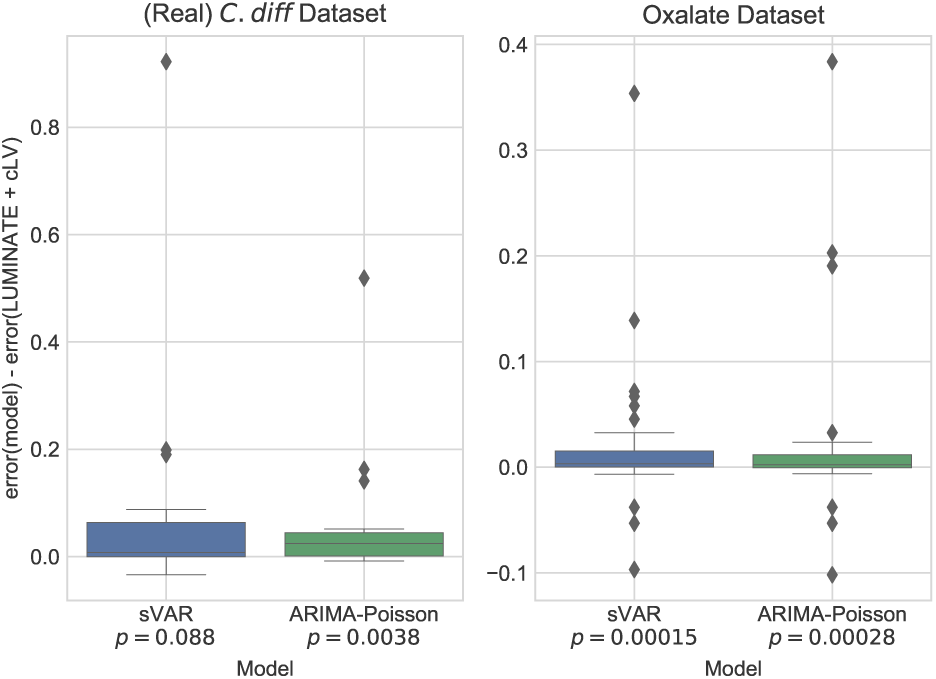
LUMINATE improves down-stream analysis. Comparison of observed and predicted trajectories between cLV fit using LUMINATE and two other models. The *y*-axis displays the difference in error from each model (*x*-axis) to LUMINATE + cLV. Significance is computed using the Wilcoxon signed-rank test.

## 4 Discussion

Recent focus on dynamic changes in microbial communities has highlighted the importance of longitudinal modeling and data collection. Thus, there is an increasing need for methods for analyzing longitudinal data that are capable of scaling to large datasets spanning many taxa. With these goals in mind, we have proposed LUMINATE: a method for estimating relative abundances, and differentiating biological from technical zeros, in longitudinal datasets. We demonstrated that LUMINATE runs orders of magnitude faster than the current state of the art without loss of accuracy, can accurately detect biological zeros, and has utility as a preprocessing step for downstream analysis such as fitting the parameters of a dynamical system.

Though we emphasized variational inference as a tool to speed up computation, we note that this is not the only approach. In particular, Silverman et al. ^23^ propose an efficient algorithm for posterior inference in models they call marginally latent matrix-t processes, of which MALLARDs are a special case. However, there is currently no public implementation of MALLARDs in their framework. Still, MALLARDs do not distinguish biological from technical zeros, a major advantage of the present work.

There are several promising areas for future work. The true zero detection framework can be extended to include external perturbations, such as antibiotics, to assess how external factors affect risk of colonization by pathogenic bacteria. We can further expand our downstream analysis to learn biological interaction networks among taxa.

